# Brain transplantation of genetically corrected Sanfilippo B Neural Stem Cells induces partial cross-correction of the disease

**DOI:** 10.1101/2022.06.30.498131

**Authors:** Yewande Pearse, Don Clarke, Shih-hsin Kan, Steven Q. Le, Valentina Sanghez, Anna Luzzi, Ivy Pham, Lina R. Nih, Jonathan D. Cooper, Patricia I. Dickson, Michelina Iacovino

**Affiliations:** Department of Pediatrics, the Lundquist Institute for Biomedical Innovation at Harbor-UCLA Medical Center, Torrance, CA 90502, USA; Department of Neurology, the Lundquist Institute for Biomedical Innovation at Harbor-UCLA Medical Center, Torrance, CA 90502, USA; Department of Neurology, David Geffen School of Medicine at UCLA, Los Angeles CA 90095, USA; Department of Pediatrics, Washington University, Saint Louis, MO 63110, USA; Department of Pediatrics, David Geffen School of Medicine at UCLA, Los Angeles CA 90095, USA

## Abstract

Sanfilippo syndrome type B (Mucopolysaccharidosis type IIIB or MPS IIIB) is a recessive genetic disorder that severely affects the brain due to a deficiency in the enzyme α-*N*-acetylglucosaminidase (NAGLU), leading to intralysosomal accumulation of partially degraded heparan sulfate. There are no effective treatments for this disorder. In this project, we carried out an *ex vivo* lentiviral correction of neural stem cells derived from *Naglu*^*-/-*^ mice (iNSCs) using a modified enzyme in which the NAGLU is fused to an Insulin-like Growth Factor II receptor (IGFIIR) binding peptide in order to improve the cross-correction efficiency. After brain transplantation of these corrected iNSCs into *Naglu*^*-/-*^ mice and long-term evaluation of the cross-correction, we successfully detected NAGLU-IGFII activity in all transplanted animals, as well as decreased lysosomal accumulation and reduced astrocytic and microglial activation throughout the transplanted brain. In addition, we identified a novel neuropathological phenotype in untreated brains characterized by decreased levels of MAP2 protein and accumulation of synaptophysin-positive aggregates in the brain. Following transplantation, this *Naglu*^*-/-*^ -specific phenotype was altered with restored levels of MAP2 expression and significantly reduced formation of synaptophysin-positive aggregates. Our results demonstrate the feasibility and long-term benefit of genetically corrected iNSCs transplantation in the Sanfilippo B brain and effective cross-correction of Sanfilippo-associated pathology in *Naglu*^*-/-*^ mice. Our findings suggest that genetically engineered iNSCs can be used to effectively deliver the missing enzyme to the brain and treat Sanfilippo type B-associated neuropathology.

## INTRODUCTION

Sanfilippo syndrome type B (Mucopolysaccharidosis type IIIB, MPS IIIB), a sub-type of Sanfilippo syndrome, is a rare inherited lysosomal storage disease caused by the deficiency of the enzyme α–*N*-acetylglucosaminidase (EC 3.2.1.50 (NAGLU)), a lysosomal acid hydrolase involved in the breakdown of heparan sulfate (HS)^1-3^. HS glycosaminoglycans (GAGs) support multiple essential roles in cellular homeostasis such as ligand binding, cellular processes, and signalling pathways activation^4-6^. Inherited mutations of the *NAGLU* gene result in a marked reduction of its enzymatic activity and the subsequent accumulation of HS-GAGs and monosialic gangliosides (GM_2_ and GM_3_) within lysosomes and other cellular compartments^7-9^. This abnormal accumulation is thought to directly impair autophagy^10^ and endosomal trafficking^11^, leading to an excessive inflammatory response affecting neuronal function and viability^12-14^.

Sanfilippo type B symptoms typically present in early childhood in the form of severe neurocognitive decline and progressive loss of speech and mobility, often leading to premature death before reaching adulthood^1,15^. There are currently no effective treatments for Sanfilippo type B patients, beyond symptomatic or palliative approaches^16^. To date, intra-cerebroventricular (ICV) enzyme replacement therapy (ERT)^17^, gene therapy^18,19^, hematopoietic stem cell gene therapy^20^, and substrate reduction^21^ have all been explored as potential treatments for this disease. However, although these approaches have shown therapeutic benefit in predominantly somatic and non-neuropathic forms of MPS, no clinical trial have succedeed in restoring or delaying MPS-associated neurological decline due to unsuccessful delivery of the therapeutics to the brain. Although systemic ERT has shown promise in promoting functional recovery in MPS subtypes known to induce moderate neurological impairment (MPS I^22^, II^23^, IVA^24^ and VII^25^), it poorly impacted both neurological deficit and cellular uptake in Sanfilippo type B because the recombinant human NAGLU (rhNAGLU) does not cross the blood-brain barrier^26,27^. Currently, there is one ERT-based clinical trial (NCT03784287) testing the effect of the ICV implantation of a NAGLU reservoir for direct delivery of the enzyme to the brain. This fusion protein consists of an IGFII receptor-binding peptide linked to the C-terminal of rhNAGLU in order to improve its cellular uptake via IGFII binding sites on the mannose-6-phosphate receptor (M6PR)^28,17^ in a glycosylation-independent lysosomal targeting (GILT) fashion^28,29^.

Gene therapy represents a promising direct and long-term approach for the treatment of single-gene disorders such as Sanfilippo type B ^19,30,31^. To date, two distinct human clinical trials using adeno-associated viruses (AAVs) have been conducted using 1) a rAAV2/5-hNAGLU vector (NCT03300453), which showed amelioration of the disease^32^, and 2) a rAAV2/9.CMV.hNAGLU vector (NCT03315182) still recruiting patients. However, pre-existing immunity to AAVs is prevalent in humans and can profoundly impact the transduction efficiency^33^.

Hematopoietic stem cell (HSC) transplantation is available as a treatment for MPS I patients with neurological symptoms. Although this approach has not been approved for Sanfilippo type A or B^34^, a Phase II clinical trial is currently on-going in MPSIIIA patients (NCT04201405). The direct implantation of neuronal stem cells (NSCs) into the CNS provides an alternative approach for sustained enzyme delivery to the brain and reduction of lysosomal storage, as shown in Sanfilippo type B and MPS VII mice^35,36 37^ as well as in other lysosomal storage disorders^38-40^. Recently this approach has also shown regenerative and therapeutic potential in neurodegenerative disorders such as Alzheimer disease ^41-45^. NSCs can be generated from induced pluripotent stem cells (iPSCs), allowing the cross-correction of patient-derived own iPSCs for an optimal compatibility and reduced risk for immune rejection.

Previously, we reprogrammed *Naglu*^*-/-*^ mouse embryonic fibroblasts into iPSCs and corrected them *ex vivo* to overexpress the human *NAGLU* enzyme. We showed that corrected NSCs (*N-*iNSCs) improved the MPS-associated brain neuropathology in *Naglu*^*-/-*^ mice^35^. As described above, NAGLU-IGFII is under clinical investigation for ERT. In this study, we have extended our previous work by using *NAGLU*-*IGFII-*secreting NSCs (*N-IGFII-*iNSC) to determine whether *NAGLU-IGFII* can perform an effective cross-correction of the Sanfilippo type B mice neuropathology.

## RESULTS

### NSC Cell Engraftment and Cell Fate

To correct the MPS IIIB-associated genetic defects, we overexpressed full-length human *N-IGFII* cDNA in *Naglu*^*-/-*^ iNSCs (as previously described^35^), via lentiviral transduction. The viral construct carries a GFP reporter via IRES sequence. Following transduction, the cells were sorted using fluorescent activated cell sorting (FACS) to obtain pure un-silenced expression of NAGLU-IGFII (**Supp. Fig. 1**). We found that *N-IGFII*-iNSCs had a 21-fold increase in intracellular NAGLU activity (p = < 0.0001), and a 4.8-fold increase in secreted NAGLU compared with wild-type (WT) cells (p = < 0.0001) (**Supp. Fig. 1-C**). Our construct was designed to address the limitation of poorly phosphorylated recombinant NAGLU, which has previously demonstrated little to no intracellular uptake. To determine whether secreted NAGLU from *N-IGFII*-iNSCs was able to enter *Naglu*^*-/-*^ cells through mannose-6-phosphate (M6P) or IGFII-dependent signalling, we carried out cellular uptake assays using excess amount of M6P and/or IGFII to inhibit NAGLU uptake via M6P receptors (M6PR). We harvested the supernatant from *N-IGFII*-iNSCs iNSCs and applied it to *Naglu*^*-/-*^ iNSCs in the presence of 5 mM M6P and IGFII. After 4 hours of treatment, we found a 3.5-fold increase in NAGLU activity compared with NAGLU-/-levels. Intracellular uptake was partially inhibited by M6P and completely inhibited by the presence of IGFII or a combination of both (**Supp. Fig. 1**). These data provide evidence that *N-IGFII*-iNSCs are capable of cross-correcting *Naglu*^*-/-*^ cells through the secretion of appropriately phosphorylated NAGLU and M6PR-dependent signalling. To evaluate their therapeutic potential *in vivo*, we implanted corrected NSCs into the brain of newborn mice and evaluated the long-term effect (9 months) of the transplantation on the disease phenotype. We have previously reported a higher level of iNSC engraftment at 9 months compared to 2 months post-transplantation, possibly due to continuous cell proliferation over the initial period after implantation^35^. Therefore, we decided to focus this study on the longer-term effect at 9 months. First, we determined the post-transplantation fate of GFP positive iNSCs *in vivo* using immunofluorescence staining of multiple phenotypic markers for NeuN and MAP2 (neurons), GFAP (activated astrocytes), and O4 (oligodentrocytes) **(Fig. 1)**. We found that GFP positive cells engrafted in the forebrain of *Naglu*^*-/-*^ mice co-localised with NeuN, MAP2, GFAP and to a lower extent with O4, indicating that *N-IGFII-*iNSC cells retained their multipotency upon engraftment into the diseased brain, and were capable of differentiating into neurons, astrocytes, and oligodentrocytes, while some GFP positive iNSCs did not commit to either cell type.

**Figure 1.**
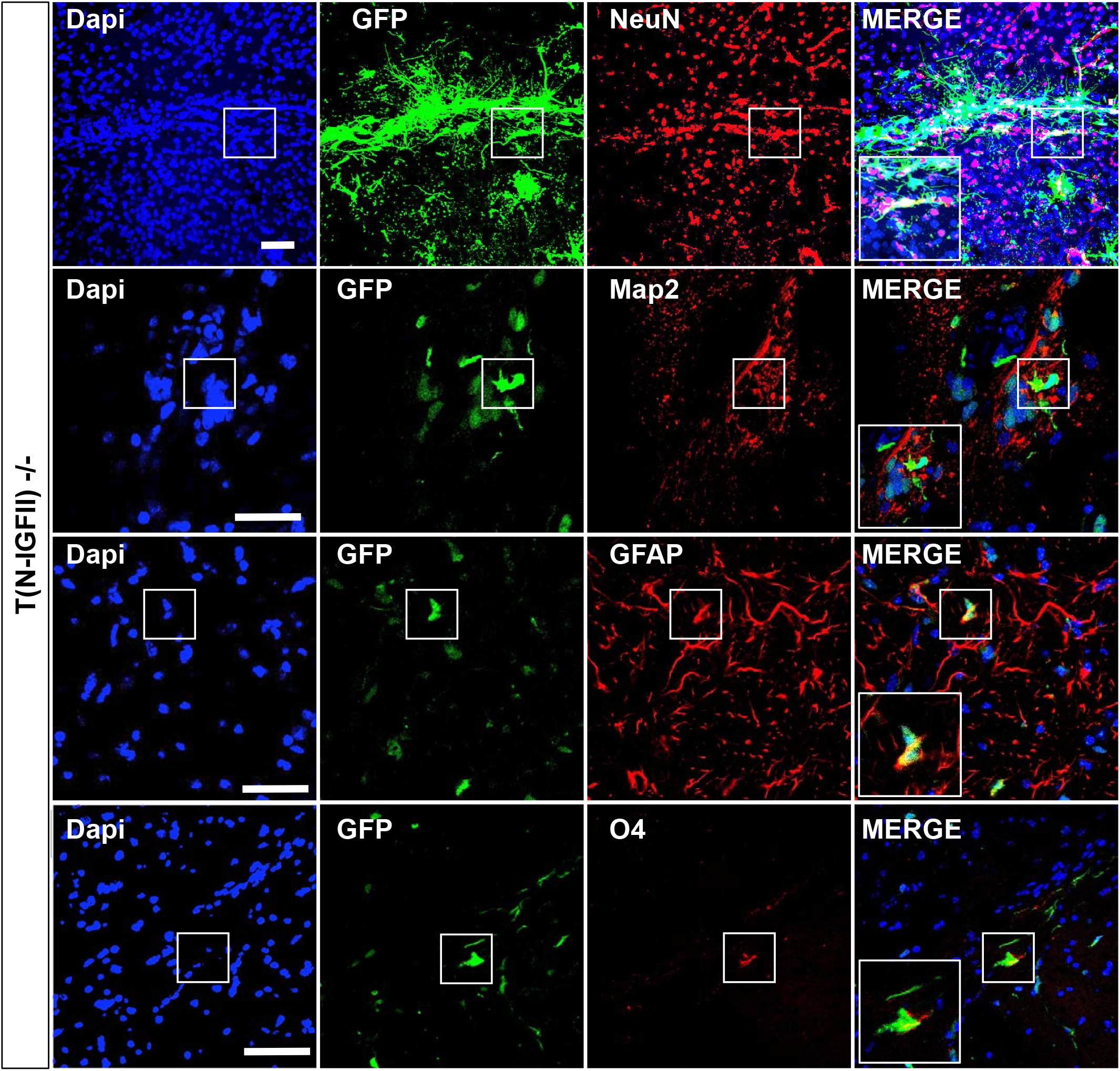
NAGLU-IGFII-corrected iNSC (*N-IGFII-iNSCs)* engraftment and cell fate 9 months after transplantation. Immunofluorescence staining of corrected *N-IGFII-*iNSCs in *Naglu*^*-/-*^ brain slices (40-μm) positive for green fluorescent protein (GFP) (green). GFP-positive *N-IGFII-*iNSCs co-localised (yellow) with glial fibrillary acidic protein (Top panel, A - D: MAP2; red), glial fibrillary acidic protein (Middle panel, E - H: GFAP; red) and oligodendrocyte marker (Bottom panel, I – L: O4; red). Nuclear staining with DAPI (blue) was overlaid. Scale bars = 50 µm.

### Pattern of NSC Cell Engraftment and N-IGFII Enzyme Activity

Next, we evaluated the engraftment of *N-IGFII*-iNSCs by measuring the percentage area of GFP staining along the rostrocaudal axis (**Fig. 2A)**. GFP positive cells were detected in all twelve mice, with a percentage area of GFP immunoreactivity measured for the whole hemisphere in each mouse. Total GFP surface ranged from 0.0001% to 25.99%, with 6 mice displaying values of 0.25% and above (0.25%, 1.16%, 1.75%, 1.75%, 2.45% 25.99%), while 6 mice showed less than 0.25% engraftment (**Fig. 2B**). The engrafted iNSCs migrated from the site of injection to spread within four brain regions along the rostrocaudal axis. To evaluate this distribution we chose 3 animals representing variable patterns of distribution regardless of their level of engraftment (**Fig. 2C)**. In animal #4763 (1.75% GFP area), the *N-IGFII-*iNSCs engrafted sporadically throughout the brain, however proportionally more cells engrafted within the caudal part of the midbrain in the substantia nigra. In contrast, animal #4871 (25.99 % GFP area) showed a high number of cells located in the olfactory bulb with gradually decreasing number towards the midbrain, with a higher number of cells detected in the substantia nigra. In animal #4872 (1.16% GFP area), higher engraftment was observed within the cerebral cortex, through the rostral portion of the hippocampus and in the substantia nigra (**Fig. 2C**). Consistent with the distribution of engrafted iNSCs, we detected NAGLU enzymatic activity in all twelve mice ranging from 0.007 to 0.492 units/mg protein (mean average enzymatic activity in the brain). Studies have shown that 1–5% of wildtype enzyme activity is sufficient to correct storage accumulation in lysosomal storage disorders (LSDs)^46,47^. In our study, all experimental animals reached 10% of carrier level (0.00775 units/mg protein red line) (**Fig. 2D)**. The overall enzyme activities correlated (Pearson R = 0.84) with the level of iNSC engraftment into the brain, as determined by assessing the GFP positive area **(Fig. 2E)**.

**Figure 2.**
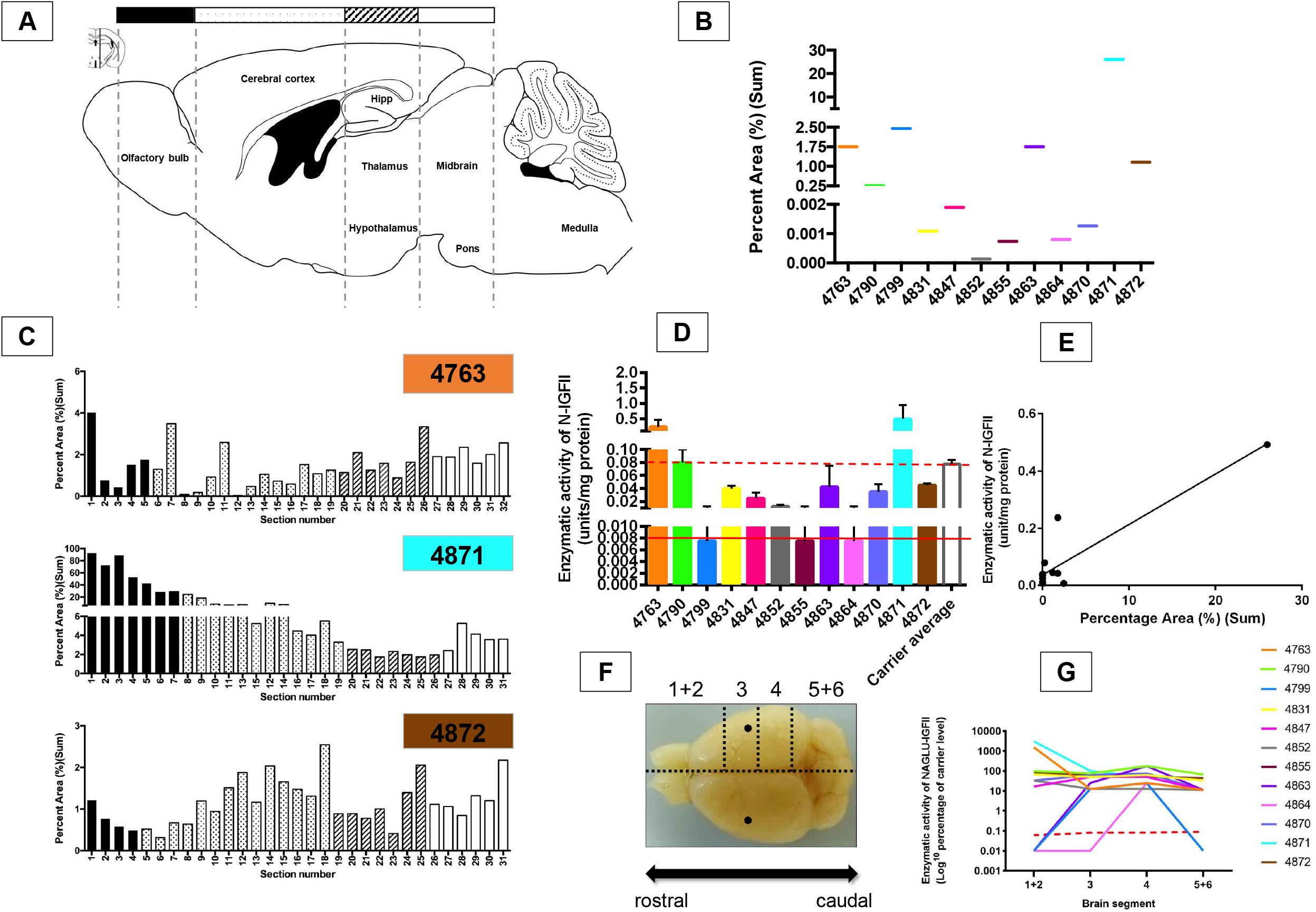
Engraftment and enzyme activity evaluation after 9 months post-transplantation. (A) Schematic illustration of a sagittal brain section (Lateral 0.24 mm Bregma) adapted from ‘The Mouse Brain in Sterotaxic Coordinates (second edition)’ by Paxinos and Franklin^77^. Black, spotted, striped and plain white bars above schematic diagram correspond to specific regions of analysis along the rostrocaudal axis shown in (C). (B) Histogram showing total percentage area (%) (sum) GFP staining in each engrafted animal plotted along the x-axis (12 animals in total). (C) Histogram showing GFP percentage area stained in three representative animals (#4763, #4871, #4872) for each individual section through the rostrocaudal axis, at the level of the olfactory bulb (black bars), cerebral cortex through to the beginning of the hippocampus (spots), thalamus (stripes) and midbrain (plain white). (D) Enzymatic activity of NAGLU-IGFII (N-IGFII) shown in units per milligram of protein, performed on brain lysates (mean average of the 4 sections depicted in Figure 5G) following injections with *N-IGFII-*iNSCs in all twelve animals, using the NAGLU enzymatic activity in *Naglu*^*-/-*^mice as a background signal. Broken red line shows protein level at 100% of the carrier level, solid line shows 10% of the carrier level (E) Correlation between level of engraftment and enzymatic activity of NAGLU-IGFII (R = 0.83) in transplanted animals. (F) Schematic diagram depicting how the brains were divided for each animal. Brains were dissected sagittally along the midline first and then right hemisphere was further sectioned into 4 slices as shown. The black dots represent the approximate intraparenchymal injection sites. (G) Graph showing the percentage (Log^10^) of carrier-level enzymatic activity of NAGLU-IGFII in 4 segments (depicted in F) along the rostrocaudal axis in twelve engrafted brains. Brain segments 1 and 2 (1+2), and 5 and 6 (5+6) were pooled. Red dotted line represents carrier level (Log^10^), solid colored lines represent each engrafted brain. Values are shown as mean ± SEM of enzyme activity measured in each slice (n = 12). Blue and purple dotted line define 10% and 100% of carrier level NAGLU-IGFII enzyme activity, respectively.

Since our analysis of GFP-positive cells revealed that *N-IGFII-*iNSCs do not distribute along the rostrocaudal axis of the brain equally, we divided the right hemisphere into 6 segments along the rostrocaudal axis, including the cerebellum and the brain stem, and measured the NAGLU enzymatic activity in each of these segments (**Fig. 2F**). Overall, the majority of injected mice displayed enzyme activity above 10% of carrier level (depicted by dotted red line) in all brain sections, and ranged from 12 – 3285.7% of the carrier control (**Fig. 2G**).

### Effect of *N-IGFII-*iNSCs on Glial Activation and Storage Accumulation

Glial activation is a characteristic feature of the neuropathology observed in both Sanfilippo type B patients and *Naglu*^-/-^ mice, and is known to intensify with age ^14,19,48,49^. To determine the impact of engrafted *N-IGFII*-iNSCs upon the activation of microglia and astrocytes as part of the innate immune response, we stained sections of 1) unaffected *Naglu*^*+/-*^ control mice (Unaffected +/-), 2) vehicle-injected *Naglu*^*-/-*^ (Vehicle -/-), and 3) *N-IGFII*-iNSC-grafted *Naglu*^*-/-*^ (T (N-IGFII)-/-) for the markers CD68 (microglial) and GFAP (astrocytes).

To determine the extent of the pathological correction, we carried out threshold image analysis of CD68, GFAP, and Lamp1 stainings along the entire rostrocaudal axis. We found a significant increase of CD68 and GFAP positive signal in the forebrain of vehicle-injected *Naglu*^*-/-*^ mice (Vehicle -/-), compared with control mice (Unaffected +/-) (**Fig. 3A, B)**, data that is consistent with our previous findings^35^. We found a 72-fold increase of the CD68 staining in vehicle-injected *mice compared with* unaffected ones (p = <0.0001). However, the brain transplantation of *N-IGFII*-iNSC was associated with a 1.84-fold reduction of CD68 signal in *Naglu*^*-/-*^ mice compared with vehicle-injected mice (p = 0.0360), and with unaffected mice (p = 0.0193, **Fig. 3A**). A similar trend was observed in GFAP signals, with a 2.2-fold increase in vehicle-treated *Naglu*^*-/-*^ mice compared with unaffected mice (p = 0.0007). Meanwhile, *N*-*IGFII-*iNSCs treatment in *Naglu*^*-/-*^ mice was associated with a reduction of GFAP signal compared with vehicle-treated mice, placing the GFAP value to comparable levels found in unaffected mice. These results indicate that *N-IGFII-*iNSCs have an overall suppressive effect on the glial reaction typically associated with Sanfilippo type B (**Fig. 3B)**.

**Figure 3.**
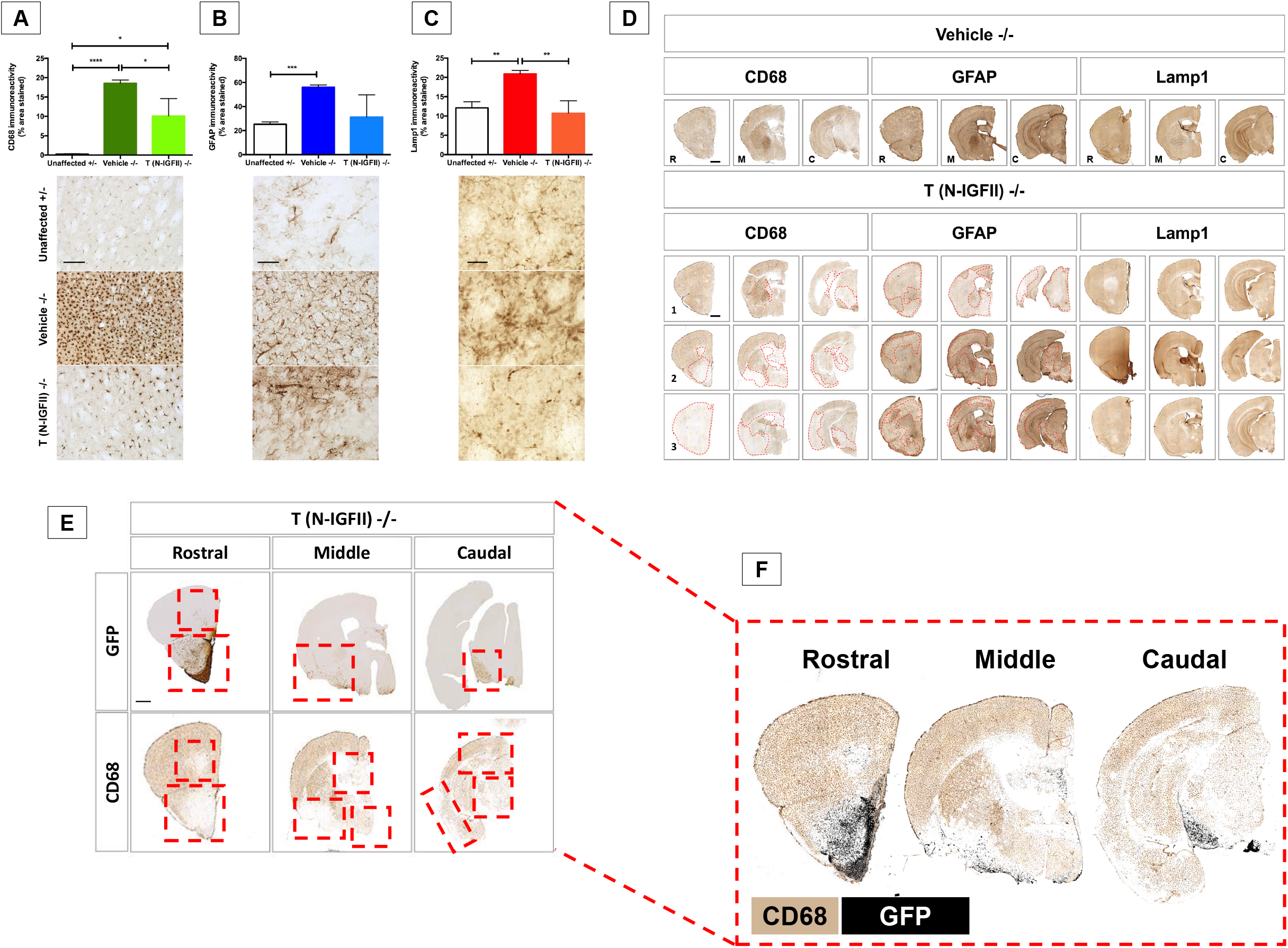
Correction of neuropathology and storage accumulation in *Naglu*^*-/-*^ mice treated with NAGLU-IGFII-corrected iNSC (*N-IGFII-iNSCs)* at 9 months. (A) Low magnification of bright-field images of coronal sections from a representative engrafted *Naglu*^*-/*^ (T (N-IGFII)-/-) mouse immunohistochemically stained for GFP, CD68 and GFAP to show the overlap between engrafted (GFP-positive) cells and pathology correction. Sections were taken at three levels along the rostrocaudal axis: rostral (at the level of the isocortex and olfactory areas), middle (at the level where the fimbria of the hippocampus appears), and caudal (at the level of the midbrain. Histograms (B–D) and representative images taken within the striatum (below) showing the percentage area of CD68 (B) and GFAP (C) immunoreactivity, and mean intensity of Lamp1 (D) immunoreactivity in every one-in-twelve sections including all regions within the forebrain, for each group measured. *p ≤ 0.05, ***p ≤ 0.001 ****p ≤ 0.0001; two-tailed, unpaired parametric t-test. Values are shown as mean ± SEM (n = 3 mice per group). (E) Representative overlapping sections from an engrafted *Naglu*^*-/-*^ mouse immunohistochemically stained for CD68 (top, color) and GFP (bottom, black and white). Sections were taken at the same levels along the rostrocaudal axis as above. Areas where CD68 staining was absent (no pixels) were modified to appear transparent, and positioned on top of the corresponding GFP-stained section (black) as an overlay (F). Low magnification of representative bright-field images of coronal sections for a representative vehicle-treated (Vehicle -/-) mice compared to three representatives N-IGFII treated (T (N-IGFII)-/-) mice to show the level of variation of pathology correction. Sections were taken at three levels along the rostrocaudal axis: rostral (R; at the level of the isocortex and olfactory areas), middle (M; at the level where the fimbria of the hippocampus appears), and caudal (C; at the level of the midbrain) of mice from engrafted *Naglu*^*-/-*^ mice, and immunohistochemically stained for CD68 (first block), GFAP (second block) and Lamp1 (third block). 1 – 3 represent three different *Naglu*^*-/-*^ mice injected with *N-IGFII-*iNSCs to demonstrate variation in CD68, GFAP and Lamp1 immunoreactivity within the same treatment group. Red broken lines outline the areas where CD68 and GFAP immunoreactivity is reduced. Post-acquisition processing was applied to all images and included adjustments to brightness and contrast and RGB curves using Adobe Photoshop CS6 to improve visibility and consistency in color tone. Scale bars = 200 µm (-D), 1000 µm (F).

Lamp1 is a lysosomal membrane protein used as a surrogate marker for lysosomal storage accumulation^50^. As expected, Lamp1 levels were significantly greater (1.7-fold) in vehicle-injected *Naglu*^*-/-*^ mice compared with unaffected controls (p = 0.0085). Strikingly, treatment with *N*-*IGFII-*iNSCs reduced Lamp1 signal to levels comparable with those found in unaffected mice (p = 0.0080, **Fig. 3C)**. In addition, correction of the inflammatory response and storage accumulation was most pronounced at the site of engraftment (the olfactory bulb, white matter tracts, the ventral forebrain, and the substantia nigra) while showing varying degrees of success beyond the site of engraftment (**Fig. 3D**).

To determine the correlation between engraftment and pathological correction, we compared side by side stainings of GFP-transplanted cells (dark color) and of CD68 microglial (**Fig. 3E-F**). We found a clear overlap between sites of engraftment and low CD68 signal (**Fig. 3F**).

### Effect of *N-IGFII*-iNSCs on Cytoskeletal Pathology and Synaptophysin Aggregation in *Naglu*^*-/-*^ Mice

We have demonstrated that grafted *N-IGFII-iNSCs* correct the glial activation typically associated with NAGLU deficiency. However, the human Sanfilippo type B syndrome is a neurodegenerative disorder that ultimately results in pronounced neuronal loss and brain atrophy^51^. We found that the extent of neuronal loss is not recapitulated in the *Naglu*^*-/-*^ mouse model^14^. In order to assess neuronal integrity, we stained for the cytoskeletal protein microtubule-associated protein 2 (MAP2), known to play an important role in stabilizing microtubules in dendritic processes^52^. Down-regulation of MAP2 has been consistently reported in the studies using neurodegenerative mouse models^53,54^. To assess whether *Naglu*^*-/-*^ mice have a similar phenotype, we stained for MAP2 in *all three mice groups* at 9 months of age (**Fig. 4A)**. We found a clear down-regulation of MAP2 signal in vehicle treated *Naglu*^*-/-*^ mice compared to unaffected mice in the hippocampus, which has intense MAP2 staining in unaffected animals (higher magnification images in red box). Treatment with *N-IGFII*-iNSC cells restored MAP2 signals to levels similar to those found in unaffected controls (**Fig. 4B**). Our quantification showed that MAP2 signal had a significant decrease in the hippocampus (p =0.0017) compared with unaffected mice. Upon *N-IGFII*-iNSC treatment, MAP2 signal was restored to unaffected level. We quantify MAP2 levels in whole brain by western analysis, but we did not detected differencences between unaffected, disease and *N-IGFII*-iNSC treated animals (**Fig. 4C**). Our data provide novel evidence that MAP2 is down-regulated in the hippocampus of *Naglu*^*-/-*^ mice. This may be associated with cytoskeletal dysfunction, but is largely prevented by *N-IGFII*-iNSC treatment.

**Figure 4.**
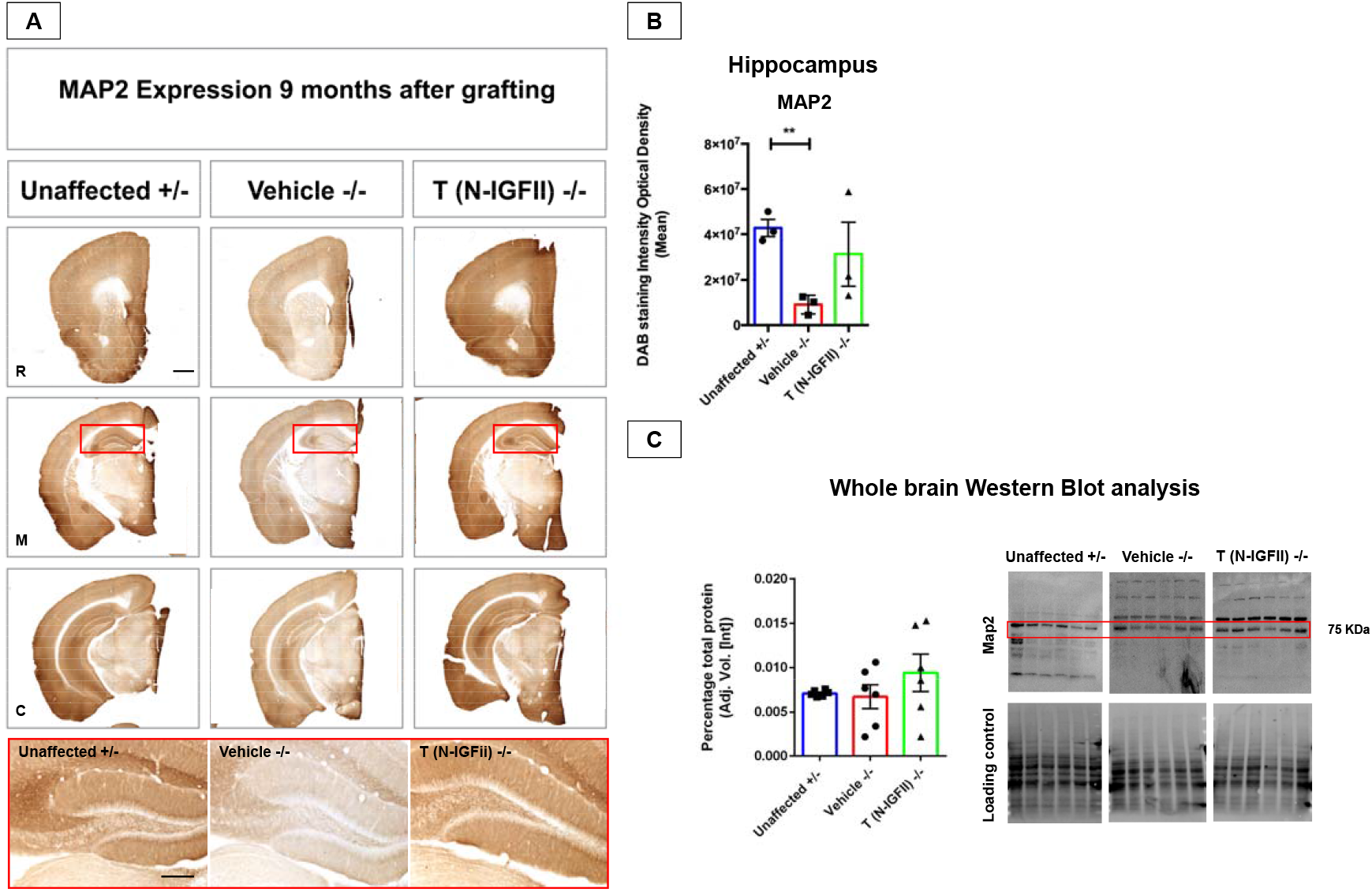
Correction of MAP2 downregulation in *Naglu*^*-/-*^ mice at 9 months. (A) Representative bright-field images of 40-μm thick half-brain coronal sections from heterozygous control mice (Unaffected) and *Naglu*^*-/-*^ mice injected with vehicle (Vehicle -/-) or *N-IGFII-*iNSCs (T (N-IGFII)-/-) immunohistochemically stained for MAP2. Panels depicted by R, M, C, show sections taken at three levels along the rostrocaudal axis: rostral (R; at the level of the isocortex and olfactory areas), middle (M; at the level where the fimbria of the hippocampus appears), and caudal (C; at the level of the midbrain). Panel highlighted in red box below, shows high magnification images of hippocampus depicted for each group. Post-acquisition processing was applied equally to all images and included adjustments to brightness and contrast and RGB curves using Adobe Photoshop CS6 to improve visibility and consistency in color tone. Scale bars = 1000 µm (top) 200 µm (bottom). (B) Histograms show the optical density (DAB staining intensity Optical Density) of MAP2 immunohistochemical staining in the hippocampus, of animal groups in (A). **p ≤ 0.01, two-tailed, unpaired parametric t-test. Values are shown as mean ± SEM (n = 3 mice per group). (C) MAP2 levels in whole-brain (n=6 for each group) measured by western blot and quantification on animal groups in (A).

Synaptophysin is an abundant integral membrane glycoprotein that is found in presynaptic vesicles of neurons. It has been previously reported that the accumulation of synaptophysin in the corpus callosum is a reliable marker of axonal damage in inflammatory conditions^55^. We found a significant increase of synaptophysin signal in vehicle-treated *Naglu*^*-/-*^ mice compared with both unaffected control mice and *N-IGFII*-iNSC-treated mice. The synaptophysin aggregates were most noticeable in the hippocampus, cortex, striatum and entorhinal cortex (EC) (**Fig. 5A**). We established a threshold level of synaptophysin signal based on these darkly stained spheroids in order to calculate the percentage area (sum) of synaptophysin-positive aggregates. Synaptophysin immunoreactivity was significantly greater (3-fold) in vehicle-injected *Naglu*^*-/-*^ mice compared with unaffected controls (p = 0.0037). Interestingly, treatment with *N*-*IGFII-*iNSCs reduced synaptophysin signal to similar levels observed in unaffected carrier mice, leading to a significant difference with values from vehicle-treated *Naglu*^*-/-*^ mice (p = 0.0146, **Fig. 5B)**. Next, we evaluated whether synaptophysin aggregates accumulate within the lysosome. We found that large synaptophysin aggregates co-localized with Lamp1 staining (**Fig. 5C**, white arrows), suggesting a lysosomal-mediated synaptophysin accumulation and lysosomal-mediated mechanism of synaptic failure^56^.

**Figure 5.**
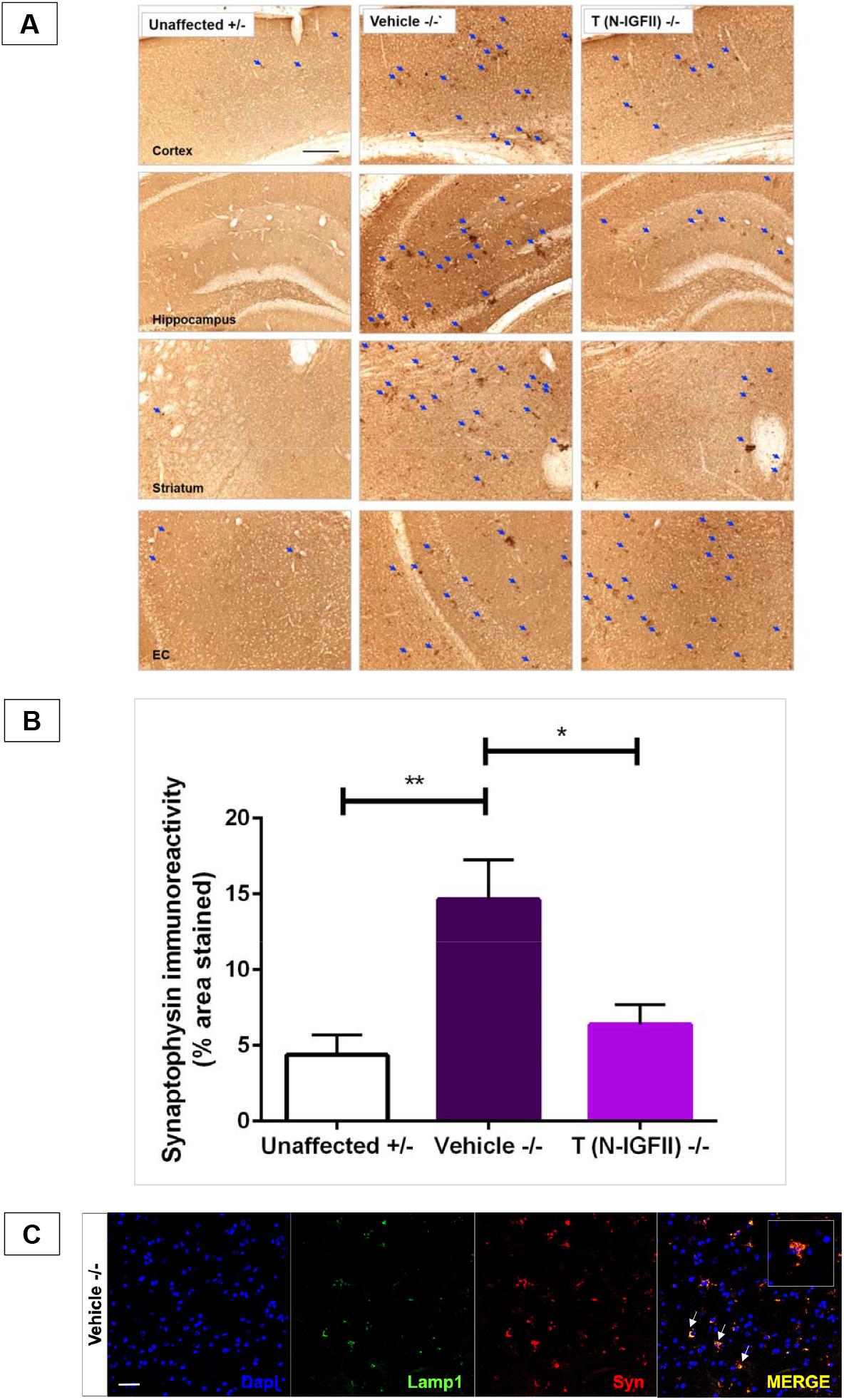
Correction of synaptophysin aggregation in *Naglu*^*-/-*^ mice at 9 months. (A) Representative bright-field images of 40-μm thick half-brain coronal sections from *Naglu*^*+/-*^ mice (Unaffected) and *Naglu*^*-/-*^ mice injected with saline (Vehicle -/-) or *N-IGFII-*iNSCs (T (N-IGFII) -/-) immunohistochemically stained for synaptophysin. Top panel shows the middle (at the level where the fimbria of the hippocampus appears) section. High magnification images, panel below, show synaptophysin-positive aggregates (blue arrows) in the cortex, hippocampus, striatum, and entorhinal cortex (EC) of all treatment groups. Post-acquisition processing was applied equally to all images and included adjustments to brightness and contrast and RGB curves using Adobe Photoshop CS6 to improve visibility and consistency in color tone. Scale bars = 1000 µm low magnification, 200 µm higher magnification. (B) Histograms showing the percentage area of Synaptophysin immunoreactivity in every one-in-twelve sections including all regions within the forebrain, for each group measured. *p ≤ 0.0146, **p ≤ 0.0037; two-tailed, unpaired parametric t-test. Values are shown as mean ± SEM (n = 3 mice per group). (C) Immunofluorescence staining showing co-localization (yellow) of Lamp1 (green) and synaptophysin (red) in 40-μm thick half-brain coronal sections from *Naglu*^*-/-*^ mice injected with vehicle (Vehicle -/-). Scale bars = 50 µm.

Together, our data show an overall reduction of synaptophysin aggregates in *N-IGFII*-iNSC engrafted mice compared with vehicle-injected *Naglu*^*-/-*^ mice. These results reveal a novel neuronal phenotype in *Naglu*^*-/-*^ brains in which synaptophysin-positive aggregates alongside Lamp1 represent a marker of neuropathological outcome that can be used to evaluate the therapeutic efficacy of engrafted corrected-iNSCs.

## DISCUSSION

Sanfilippo type B is caused by a deficiency in the enzyme NAGLU, known to be involved in the breakdown of HS, resulting in the accumulation of partially degraded GAGs in the lysosome. Pathological abnormalities in *Naglu*^*-/-*^ brains provide a way to study the disease mechanism observed in Sanfilippo patients as *Naglu*^*-/-*^ mice display abnormal lysosomal pathology, a pronounced neuroimmune response, and severe neuronal dysfunction^13,57-59.^ Previously, we showed that NAGLU produced *in vitro* by our *N*-iNSCs can be endocytosed by NAGLU-deficient cells via the M6PR and cross-correct MPS-associated microglial activation, astrocytosis, and lysosomal storage accumulation *in vivo*^35^. The current study shows that cross-correction was achieved using a modified NAGLU-IGFII enzyme. However, despite the use of NAGLU-IGFII, cross-correction was not sufficient to completely correct the microglial activation. This result implies that limited cross-correction (due to insufficient enzyme distribution) might require additional soluble factors from iNSCs to dampen inflammation. Therefore, widespread dissemination of iNSCs or a combinatorial approach using an anti-inflammatory drug may prove more effective.

Previous studies on the post-transplantation fate of iNSCs in the adult brain have focused on direct migration toward focal areas of injury e.g. ischemia or brain tumors^60,61.^ However, little is known about the way NSCs migrate in un-lesioned brain regions following intraparenchymal transplantation. Although transplanted *N-IGFII*-iNSCs were detected throughout the rostro-caudal axis in the majority of transplanted brains 9 months after treatment, their pattern of engraftment was highly variable, which can be attributed to the intrinsic characteristics of the donor cells. Overall, *N-IGFII-*iNSCs were found mostly within the ventral forebrain and substantia nigra (**Fig. 2**). Future studies are needed in order to evaluate the normal migratory behaviour of transplanted *N-IGFII-*iNSCs in the brain of wild-type neonatal mice. In addition, additional safety studies are required to assess the proliferation rate and cell death rate of donor cells in transplanted mice.

MAP2 is a cytoskeletal protein found in the neuronal dendritic compartment where it plays an important role in the morphological stabilization of dendritic processes. It is a marker of structural integrity and its expression has been linked to dendritic outgrowth, branching, and post-lesion remodelling and plasticity^62^, known to play an important role in memory in the hippocampus^63^. Down-regulation of MAP2 expression has been linked with impaired microtubule assembly and has been found in the hippocampus of the aging rat brain and Alzheimer disease^64-66^. In our study, a reduced MAP2 expression was specifically found within the hippocampus of *Naglu*^*-/-*^ mice. Severe dementia is one of the primary characteristics of Sanfilippo type B disease in humans, and *Naglu*^*-/-*^ mice perform poorly in water maze and radial-arm maze behavioural tests indicating an impairment in spatial learning^67^. Therefore, MAP2 down-regulation may partially explain the dementia-like symptoms reported in Sanfilippo type B disease^1,67^.

Aggregation of abnormal cellular proteins into insoluble complexes is involved in the pathogenesis of most human neurodegenerative diseases^68^. Partially degraded HS is the primary storage product in NAGLU-deficient lysosomes. In addition, NAGLU-deficient neurons were found to accumulate GM3 ganglioside and cholestero^l9,69^, ubiquitin, and subunit c of mitochondrial ATP synthase (SCMAS)^70^. However, the mechanism by which abnormal protein aggregation leads to the degradation of different neuronal populations is not fully understood. In this study, we identified aggregates of synaptophysin in the cells of *Naglu*^*-/-*^ mice. Synaptophysin aggregation has been described as a reliable marker for axonal damage in conditions such as multiple sclerosis, CNS trauma, and multiple neuroinflammatory conditions^55^. Decreased synaptophysin expression has been reported in several models of neurodegenerative disease^71-73^. A previous study has demonstrated that synaptophysin expression is decreased in Sanfilippo type B as a result of storage material accumulation, and can be reversed following correction by gene transfer. Several studies have reported that synaptophysin levels were reduced in the rostral cortex of Sanfilippo type B mice compared to controls as early as 10 days and that this reduction could not be accounted for by neuron loss^14,74^. In addition, published literature has showed that synaptophysin does not accumulate in the lysosome and that dysregulation of synaptophysin levels is a result of enhanced proteasomal degradation induced by GAGs^14,74^. In our study, we found that large synaptophysin positive aggregates were present in multiple brain regions, which could be explained by the impaired degradation of the protein, despite a down-regulation of its gene expression. Our thresholding analysis methodology was not able to detect the synaptophysin down-regulation in Sanfilippo type B brains, instead we report accumulation in lysosomes. Studies suggest that the formation of such cytoplasmic inclusion bodies requires active, retrograde transport of aggregates, or misfolded protein on microtubules. A microtubule-based apparatus, the aggresome, forms and isolates protein aggregates within the cytoplasm^47,75^. Our MAP2 and synaptophysin data suggest that the down-regulation of MAP2 observed in our study may be indicative of microtubule dysfunction that results in the aggregation of synaptophysin.

We previously transplanted mice with NAGLU-expressing iNSC^35^. In our initial study, we assessed engraftment and distribution at 2 and 9 months post-transplantation and detected more engrafted cells and a broader dispersion throughout the brain at the 9 months time point, most likely due to higher proliferation over the course of weeks. In addition, we found that at this time point, almost complete neuropathology correction was observed in some brain regions of low engraftment, observation from which we concluded that the corrective capability of engrafted cells relies on the total level of engraftment throughout the brain rather than the precise distribution of the cells. One strong argument is found in the correlation between total cell engraftment and amount of secreted enzyme that is then available to cross-correct neighbouring cells. Our approach using a modified NAGLU-IGFII enzyme reveals that this fusion protein can be secreted and uptake but the efficiency of cross-correction is limited by the engraftment and dissemination of NSCs in the postnatal brain. While 1-5% of enzymatic activity may be sufficient to prevent storage accumulation (**Fig. 3C**), it appears that much higher enzymatic activity is needed to counteract the MPS-associated neuroinflammation throughout the whole brain, and despite the use of NAGLU-IGFII, cross-correction was not sufficient to improve the efficacy of this treatment. As shown in Figure 1, the GFP signal was weak, suggesting that it may be necessary to improve overexpression of NAGLU using a more powerful promoter, together with increased cross-correction via modified NAGLU enzyme.

Despite *N-IGFII-*iNSC engrafted mice having a low level of chimerism, grafted cells distributed along the entire rostral-caudal axis and were capable of providing comparable levels of NAGLU activity to those found in unaffected brains. This finding suggests that NAGLU-IGFII was sufficiently secreted and distributed widely to cross-correct the neuropathology. This cross-correction was also reflected in the systematic correction of glial activation, storage accumulation, neuropathology of the microtubule system and synaptic vesicles. In conclusion, we have shown that the successful engraftment of NSCs expressing a modified NAGLU protein can be achieved, and resolve the accumulation of storage material and glial activation in 9 month post-treated *Naglu*^*-/-*^ mice. We have also identified novel targets of neuronal pathology that can be used to more rigorously assess the efficacy of potential treatments.

## MATERIAL AND METHODS

### Lentivirus Production

The human *NAGLU* gene was engineered from a previously published *NAGLU-IGFII*^*50*^ sequence and further cloned into a lentiviral backbone plasmid, pSAM-GFP^35^ at the *XhoI* and *NotI* sites. The resulting expression vector was named pSAM-NAGLU-IGFII-IRES-GFP. The lentiviral supernatant was produced with 293T cells (American Type Culture Collection, Manassas, Virginia) as described previously^76^.

### Cellular Reprogramming

*Naglu*^*-/-*^ MEFs were transduced with the reprogramming factors pMXs-*OCT3/4*, pMXs-*SOX2*, pMXs-*KLF4*, and pMXs-c-*MYC* (Addgene, Cambridge, MA) as reported previously^77^. MEFs were cultured until individual iPSC colonies were formed. Colonies were picked and grown individually on a feeder layer of MEFs treated with mitomycin C (Roche, Indianapolis, IN) as described previously^76^. Characterization was performed using alkaline phosphatase staining (Cell Biolabs, San Diego, CA) and immunofluorescence for the presence of Oct4.

### Directed Differentiation of iPSCs to iNSCs

iPSCs were grown feeder-free at a density of 8.0 × 10^4^ cells/well in embryonic stem cell (ESC) medium (knockout DMEM (Gibco, Carlsberg, CA) with 10% fetal bovine serum (FBS) (R&D systems, Minneapolis, MN) and 500 U/mL leukemia inhibitory factor; Millipore, Burlington, MA). The following day, the medium was changed to NSC medium (DMEM/F12 supplemented with B27 without vitamin A and N2; Gibco, Carlsbad, CA) and grown for 12 days. On day 12, NSC medium, 20 ng/mL of epidermal growth factor (EGF), and 20 ng/mL basic fibroblast growth factor (bFGF) were added (PeproTech, Rocky Hill, NJ). Cells were expanded in NSC medium + EGF + bFGF as monolayers^78^.

### Enzyme Activity Assays

The catalytic activity of NAGLU was determined by hydrolysis of the fluorogenic substrate 4-methylumbelliferyl-*N*-acetyl-α-glucosaminide (EMD Millipore, Burlington, MA) with a final concentration of 0.1 mM substrate in the incubation mixture as described previously^50^. A unit of activity is defined as the release of 1 nmol of 4-methylumbelliferone (4MU) per hour at 37°C. Naglu activity signal in Naglu-/-cell and in Naglu-/-brain, was used as background and subctracted from all other measurements. Protein concentration was estimated by the Bradford method, using BSA (Bio-Rad, Hercules, CA) as the standard. Fluorescence was measured on a microplate reader (Spectramax Paradigm, Molecular Devices, San Jose, CA) with λ_ex_ 365 nm and λ_em_ 445 nm. Intracellular NAGLU activity is presented as units per milligram of protein.

### Experimental Animals

Animal experiments were approved by the Institutional Animal Care and Use Committee at the Lundquist Institute at Harbor-UCLA Medical Center, which is accredited by the Association for Assessment and Accreditation of Laboratory Animal Care (AAALAC). The *Naglu*^*-/-*^ knockout mouse was a gift from Dr. Neufeld from the University of California, Los Angeles (UCLA) was back-bred onto C57BL6/J ^79^ and maintained in Lundquist Institute. Genotyping was performed with the following primers: NAG5′, 5′-TGGACCTGTTTGCTGAAAGC-3′; NAG3′, 5′-CAGGCCATCAAATCTGGTAC-3′; Neo5′, 5′-TGGGATCGGCCATTGAACAA-3′; and Neo3′, 5′-CCTTGAGCCTGGCGAACAGT-3′. *Naglu*^*+/-*^ females were crossed with *Naglu*^*-/-*^ males to obtain homozygous affected mice and heterozygous (unaffected) controls. Experiments were performed on age-matched mice (usually littermates) of either gender.

### Injection of NSCs

2.5 × 10^5^ NAGLU-IGFII-overexpressing NSCs (*N*-*IGFII-*iNSC) in 3 μl of PBS or PBS alone were injected bilaterally into the striatum of postnatal day 0 (P0) or P1 *Naglu*^*-/-*^ neonatal mice under cryo-anesthesia by Hamilton syringe with 32G needles. The injection site was approximately 3.0 mm rostral from bregma, 4.0 mm laterally from the midline and at a depth of 3.0 mm as previously described^35^. Injections were performed manually by the same experimenter using Bregma and Lamda landmarks. A plastic block was attached to the end of the syringe. The length of the needle entering the brain, was limited by contact of the block with the top of the skull, thus ensuring the depth of injection was fixed and reproducible between injections.

### Tissue Harvesting, Processing, and Histological Staining

After 9 months post-transplantation, brains were dissected sagittally along the midline, and left hemispheres were post-fixed overnight at 4°C in 4% paraformaldehyde (PFA) before dehydration and cryoprotection at 4°C in a solution of 30% sucrose in Tris-buffered saline (TBS). The right hemispheres were further sectioned into 2 mm-thick coronal slices using an adult mouse brain slicer matrix (Zivic Instruments, Pittsburgh, PA) and rapidly frozen and stored at −80°C until performing NAGLU activity assay. For the fixed hemisphere, 40-μm frozen coronal sections were cut through the rostrocaudal extent of the cortical mantle (Microm HM 430 freezing microtome, Thermo Fisher Scientific, Waltham, MA). Sections were collected in a cryoprotectant solution (30% ethylene glycol/15% sucrose/0.05% sodium azide in TBS) and stored at 4°C before histological processing^80-82^.

### Western blotting

Protein samples and dual plus molecular weight ladders were separated by SDS-PAGE and loaded on a Precast stain free gels with a 4–15% gradient (Bio-Rad). Proteins were transferred to PVDF membranes (Bio-Rad) using the Bio-Rad Trans-Blot Turbo Transfer System. Total proteins on membranes were detected using the Geldoc system (Biorad) gel free staining. Membranes were blocked with 5% non-fat milk in TBS-T and incubated with primary antibody rabbit Anti-MAP2 (1:1000 dilution, Abcam, Cambridge, MA, catalog no. ab183830). Anti Rabbit-HRP (Invitrogen G-21234) was used as a secondary antibody. Membrane were analyzed using ChemiDoc MP (Biorad).

### Immunohistochemistry

To determine the identity of engrafted stem cells, single free-floating frozen sections were immunohistochemically stained using the following antibodies (all from Invitrogen Carlsbad, CA except specify: Rabbit anti-GFP (1:10,000 dilution, catalog no. A10259); Rat anti-GFAP (1:400 dilution, catalog no. 13-0300). Secondary detection was performed with: Streptavidin DyLight 488 nm (1:200 dilution, Vector Laboratories, Burlingame, CA, catalog no. SA-5488), Goat anti–Rat Alexa 546 nm (catalog no. A-11081), and Goat anti-Chicken Alexa 633 nm (catalog no. A-11040). All secondary antibodies were used at 1:500 dilution, with the exception of Streptavidin DyLight 488 nm. To examine the level of stem cell engraftment, the extent of glial activation, storage material accumulation and neuronal markers, adjacent one-in-six (GFP) and one-in-twelve (GFAP, CD68, Lamp1, MAP2, synaptophysin) series of free-floating frozen sections were immunohistochemically stained using a standard immunoperoxidase staining protocol, described previously^80-82^. GFP (as above), polyclonal rabbit anti-GFAP (1:8,000 dilution, Agilent Technologies, Santa Clara, CA; catalog no. Z0334), rat anti-CD68 (1:2,000, Bio-Rad, Hercules, CA; catalog no. MCA1957), rat anti-Lamp1 (1:4,000, DSHB, Iowa City, IA; catalog no. 1D4B), rabbit Anti-MAP2 (1:40,000 dilution, Abcam, Cambridge, MA, catalog no. ab183830) and Rabbit anti-synaptophysin (1:2,000 dilution, Abcam, Cambridge, UK, catalog no. ab14692) were used, followed by the appropriate secondary antisera (all from Vector Laboratories, Burlingame, CA, except specify): biotinylated goat anti-rabbit IgG (catalog no. BA-1000), biotinylated swine anti-rabbit IgG (GFAP, Agilent Technologies, Santa Clara, CA; catalog no. E0353), biotinylated rabbit anti-rat IgG (CD68, catalog no. BA-4001), and biotinylated goat anti-rat IgG (Lamp1, catalog no. BA-9400), all 1:1,000 dilution. Sections were incubated in an avidin-biotin-peroxidase complex kit (Vectastain Elite ABC Kit, Vector Laboratories, Burlingame, CA, catalog no. PK-6100). Immunoreactivity was visualized by incubation in 0.05% 3’, 3’-diaminobenzidine (DAB; Sigma-Aldrich, St. Louis, MO) and 0.01% H_2_O_2_ in TBS.

### Quantitative Analysis

To analyze the percentage area of engraftment (GFP/CD68/GFAP/Lamp1) all stained sections were scanned as follow: slides were imaged field-by-field at a magnification of x 10 (GFP, CD68 and GFAP) and x 20 (Lamp1), using a Zeiss Axio Imager M2 microscope and an automated stage (MBF Bioscience), then stitched using *StereoInvestigator* software (MBF Bioscience, Williston, VT). Thresholding analysis was performed with *Image Pro-Premier* software (Media Cybernetics, Rockville, MD) to report percentage area staining (sum) as described previously^80,83.^ The Optical Density (Intensity) was reported for MAP2 and was defined by two intensity values where darker staining is read as opaque and no staining as transparent. The standard optical density formula, which assumes that light decays exponentially as it goes through light-transmitting material (OD(x,y) = −log ((I(x,y) - BL) / (IL - BL)), where BL and IL are the Black Level and the Incident Level respectively, and I(x,y) is the intensity at location x,y) can be used to create an intensity curve where the darkest staining (black pixels) equal 0 and the lightest staining (white pixels) equals 255. The regions of interest were defined according to neuroanatomical landmarks described by Paxinos and Franklin^84^. For each sample, the appropriate threshold was applied to capture the different staining intensities for each of the areas stained (Optical Density).

Each section was outlined as a region of interest and analysed together as one whole. For each sample, the appropriate threshold was applied to capture the different staining intensities for each of the areas stained according to color, background and morphology.

### Statistical Analysis

All statistical analyses were performed using Prism software (GraphPad, La Jolla, CA). A two-tailed, unpaired, parametric t-test was used when two groups were compared. Results were considered statistically significant when p < 0.05. Unless otherwise stated, statistical comparisons are between untreated *Naglu*^*-/-*^ mice and unaffected heterozygous controls and the treatment groups and untreated *Naglu*^*-/-*^ mice. One-way ANOVA was used to compare groups greater than two.

## ACKNOWLEDGMENTS

We thank members of Dr Cooper’s Pediatric Storage Disorders Laboratory for helpful discussion and their technical support. This work was supported by grants from the NINDS (1R41NS092221-0181, 1R01 NS088766, R21NS096044) and a traineeship from 5T32 GM8243-28 (to S.-h.K.).

## AUTHOR CONTRIBUTIONS

Conceptualization, M.I. and P.I.D.; Methodology, M.I., P.I.D and J.D.C.; Validation, Y.P., D.C. and S.-h.K.; Formal Analysis, Y.P., D.C. and S.-h.K.; Investigation, Y.P., D.C., S.- h.K., S.Q.L.,V.S., M.T., and N.R-D.; Resources, M.I., P.I.D and J.D.C.; Data Curation, Y.P., D.C, S.-h.K.. Writing – Original Draft, Y.P. and D.C.; Writing – Review & Editing, Y.P., M.I., P.I.D., J.D.C. and S.-h.K.; Visualization, Y.P. and D.C. S.-h.K.; Supervision, M.I., P.I.D and J.D.C.; Project Administration, M.I. and P.I.D, Funding Acquisition, P.I.D and M.I.

## FIGURE LEGENDS

***Supplementary Figure 1. NAGLU-IGFII* Gene Therapy and Subsequent Biochemical Characterization**.

(A) FACS analysis of *Naglu*^*-/-*^ Neural Stem Cells (iNSCs) (white) transduced with *NAGLU-IGFII-GFP* (black). (B, C) Quantifying secreted and intracellular NAGLU enzyme activity in iNSCs overexpressing *N-IGFII*-iNSCs. *p ≤ 0.05, ***p ≤ 0.001 ****p ≤ 0.0001; two-tailed, unpaired parametric t-test. (D) Cellular uptake/inhibition assay on iNSCs (*Naglu*^*-/-*^) treated with supernatant collected from *N-IGFII*-iNSCs at the presence or absence of 5 mM mannose-6-phosphate (M6P), n = (3) one-way ANOVA p = <0.0001. Intracellular NAGLU activity is presented as Units/mg of protein whereas the secreted NAGLU activity is as Units/ml expressed as mean ± SD, where a unit is defined as the release of 1 nmol of 4-MU per hour at 37°C. *\*\**p < 0.01, *\*\*\*\**p < 0.0001.

