## Supplementary figures and images for "Brain transplantation of genetically corrected Sanfilippo B Neural Stem Cells induces partial cross-correction of the disease"

### Supplementary Figure 1

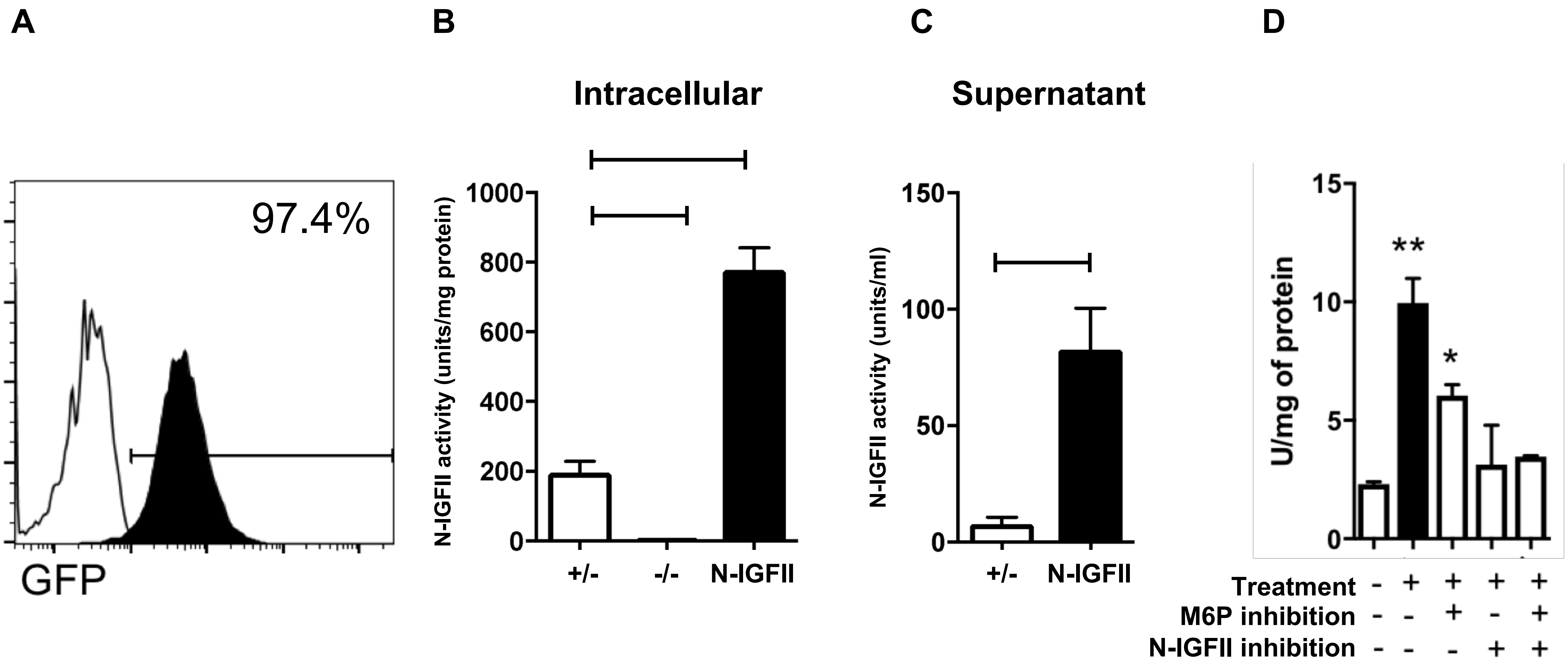
